# Dimer arrangement and monomer flattening determine actin filament formation

**DOI:** 10.1101/294256

**Authors:** Maria Hoyer, Jose Rafael Cabral Correia, Don C. Lamb, Alvaro H. Crevenna

## Abstract

Actin filament dynamics underlie key cellular processes, such as cell motility. Although actin filament elongation has been extensively studied under the past decades, the mechanism of filament nucleation remains unclear. Here, we immobilized gelsolin, a pointed-end nucleator, at the bottom of zero-mode waveguides to directly monitor the early steps of filament assembly. Our data revealed extensive dynamics and that only one, of two populations, elongates. Annalysis of the kinetics revealed a more stable trimer but a less stable tetramer in the elongating population compared to the non-elongating one. Furthermore, blocking flattening, the conformational change associated with filament formation, prevented the formation of both types of assemblies. Thus, flattening and the initial monomer arrangement determine gelsolin-mediated filament initiation.

## INTRODUCTION

Actin and actin-like proteins are ubiquitous across all phyla of life (*1*). In eukaryotes, the actin cytoskeleton is responsible for many cellular processes such as cell division, endocytosis and cell motility (*2*). This myriad of functions is intimately linked to the capacity of actin to self-associate into filaments (*2*). Actin filaments are double stranded filaments of actin monomers arranged in a head-to-tail manner (*3*). Actin monomers within the filament lattice exist in a conformation different from those free in solution; the dihedral angle formed by actin subdomains is about 20 degrees smaller in the filament (*4*) compared to the uncomplexed structure in solution (*5*). Pure actin protein molecules can spontaneously self-assembly into filaments, a process that is thought to occur via a nucleation-elongation mechanism (*6, 7*). In this model, a very fast dissociation rate (10^6^−10^8^ s^−1^) of a head-to-tail dimer and a fast dissociation of the trimer (10^3^ s^−1^) impede the formation of tetramers that, once formed, will elongate until an equilibrium is reached (*6*). The fast dissociation of dimers and trimers as well as the micro-molar amounts of actin necessary to promote filament formation have made challenging the study of the early steps during filament formation. Therefore, the kinetics of the first steps and whether conformational changes occur during nucleation remain poorly understood.

In cells, proteins called nucleators assist in the filament initiation process by interacting with actin monomers and stabilizing dimer or trimer monomer arrangements that allow filament growth (*8*). Based on the protein concentration needed to promote filament formation, nucleators are loosely defined as weak or strong (*9*). However, whether the poor nucleation ability originates from transient nucleator-actin complexes, kinetic dead ends or other possible models is not clear.

In order to shed light into some of these issues, we monitor here individual monomers during the nucleation process with the use of zero-mode waveguides ‘ZMW’. ZMW are nano-fabricated apertures on an aluminum layer deposited on a glass surface that reduces the effective observation volume to atto-liters. The reduction in the observation volume allows higher concentrations of fluorescently labeled reactants, up to micro-molar levels, to be monitored while maintaining single molecule resolution (*10*). The micro-molar amounts of labeled actin are needed to overcome the critical concentration for filament formation (*11*). We used actin protein molecules covalently labeled with atto-488 on random surface lysines, which is known not to impair filament nucleation and elongation (*12*). To help filament initiation and to anchor nascent oligomers to the surface, we used Gelsolin, a Ca^2+^-dependent ‘weak’ pointed-end nucleator (*13*). Gelsolin can simultaneously interact with two actin molecules, forming a gelsolin-(actin)_2_ ‘GA_2’_ complex (*14*). However, the GA_2_ only acts as a weak nucleator as its presence in bulk actin polymerization assays does not abolish the lag phase (*14*) compared to the activity of ‘strong’ nucleators (such as Lmod (*15*)). To investigate gelsolin-mediated actin filament formation, we immobilized full-length gelsolin molecules at the bottom of ZMW via biotin functionalization. Biotinylation of gelsolin did not alter its capacity to bind and cap growing actin filaments (Fig. S1). We placed ZMW arrays on a fluorescence microscope and monitored hundreds of individual apertures in real time. Addition of 1 µM actin-atto488 under polymerization conditions (in the presence of KCl and Ca^2+^) to the ZMW with surface-immobilized gelsolin (see Materials and Methods for details) resulted in assembly dynamics with step-wise changes of fluorescence intensity over time (Fig. 1). Without gelsolin or in the absence of KCl and Ca^2+^, no increase in intensity was observed, indicating that the increase in fluorescence signal over time was due to specific binding of actin monomers to activated gelsolin (Fig. 1, S2). To minimize the possibility that two or more molecules were present in a single aperture, we used a low concentration of gelsolin when incubating the waveguides. The fraction of apertures monitored where signal was detected was never more than 30% as required by Poissonian statistics to keep the double occupancy of apertures low. Therefore, the use of ZMW allows monitoring the self-assembly of single actin monomers as they associate with individual gelsolin or with actin molecules in real time.

**Figure 1:**
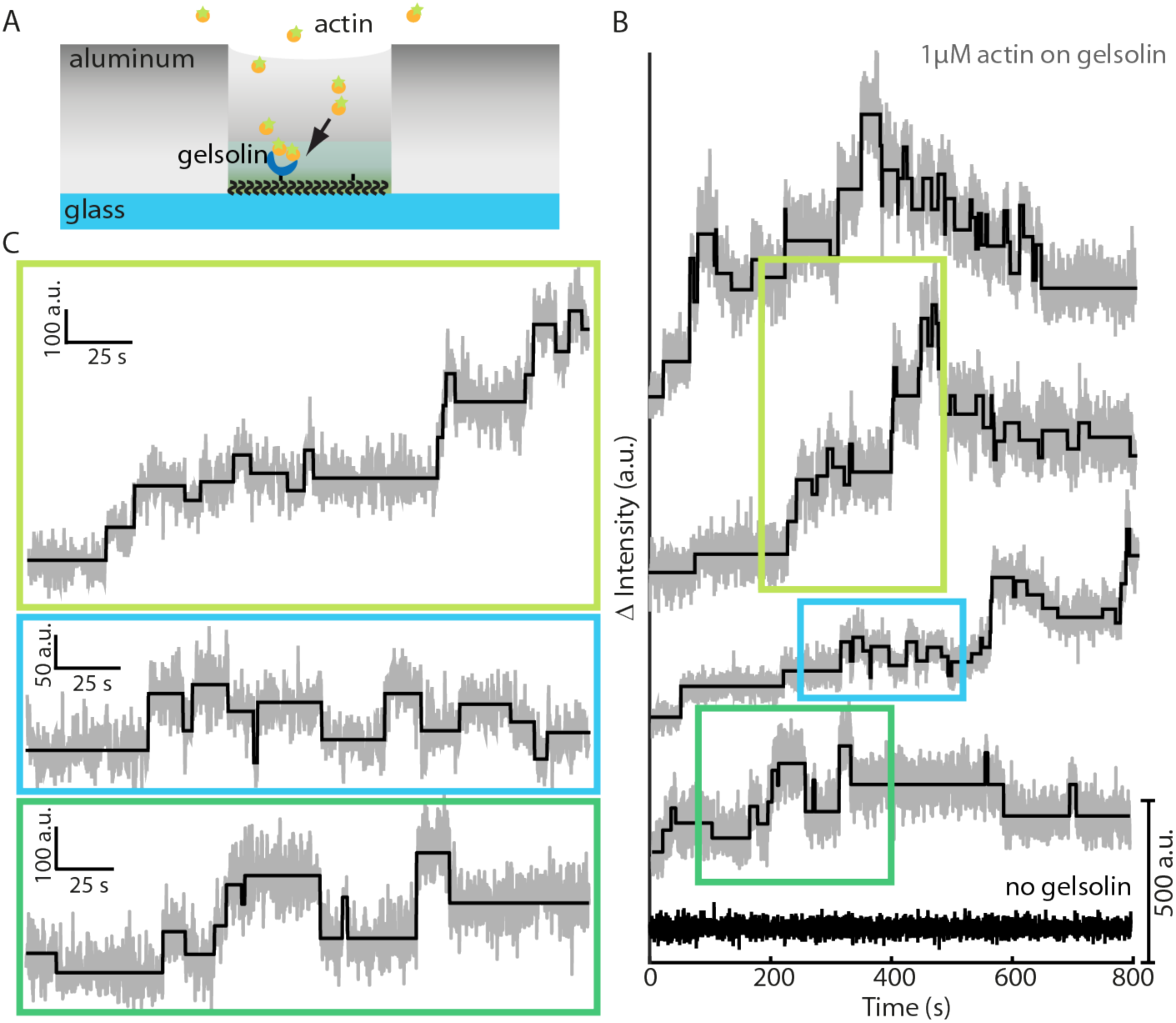
Observation of gelsolin-mediated actin filament nucleation and elongation within zero-mode waveguides (ZMW). (A) Schematic of a ZMW with gelsolin attached to the coverslip. (B) Experimental data. 1 uM of atto-488 labeled actin was added to the ZMW and the fluorescence was monitored over 14 minutes. Grey: measured intensity trace, Black: underlying trace as determined by a step-finding algorithm. The lowest traces was taken from an experiment where gelsolin was not present. (C) Zoom-ins of the corresponding highlighted regions in panel B.

In general, fluorescence intensity traces showed many up and down steps (Fig. 1, 2 and S3 for a gallery). Some traces displayed what could be considered growth while other traces, although dynamic, appeared to stagnate around a certain intensity level (Fig. 1). The concentration of actin used (1µM) is above the critical concentration for pointed-end growth (600 nM) (*16*) and, therefore, we had expected to observe mainly continuous filament elongation for all formed assemblies. In order to investigate this discrepancy in more detail we compared the expected size distribution of filaments, using literature values, with our data. We estimated the size of filaments formed on the apertures by converting the intensity level into a monomer number by first running a step-finding algorithm (*17*) (Fig. 2A left y-axis) and then applying an intensity-to-monomer conversion algorithm (see material and methods for details) (Fig. 2A right y-axis). Before applying the step finding algorithm to analyze experimentally obtained data, we tested its performance on simulations (Fig. S4) and on experimental data from molecules with a known number of fluorophores (Fig. S5). Analysis of experimental data from DNA molecules with 4, 6 or 8 fluorophores recovered well the known number of fluorophores and showed a similar intensity step size distribution to the one obtained from actin (Fig. S6). The estimated monomer number trace describes well the raw intensity trace (Fig. S7, S8). This analysis allowed us to compare the filament size in number of monomers per aperture at the end of the measurement with the expected behavior based on literature values (see Materials and Methods for details). The observed monomer number per filament at 800 s was ~5, which is much below the expectation of ~40 monomers. Overall, the observation of an elongating filament was a rare process, as only about 30% of all recorded traces had accumulated more than 6 monomers within 800 s whereas all other had 5 monomers or less (Fig. S9). Next, we investigated possible causes for the observed monomer number distribution. First, we included irreversible photobleaching as a possible source for the discrepancy (see Materials and Methods for details). Including an experimentally determined photobleaching rate of 0.003 s^−1^ (Fig. S10) into the expected filament size distribution does not reproduce the observed oligomer size distribution at 800 s (Fig. 2B). Another explanation would be that some of the surface-immobilized gelsolin has lost its capacity to bind actin monomers. We observed that the biotinylated gelsolin is functionally equivalent to its non-biotinylated form (Fig. S1) and that surface-immobilization does not preclude its interaction with actin (Fig. 1, S10). Therefore, neither photobleaching nor surface-induced artifacts can account for the observed filament size distribution.

**Figure 2:**
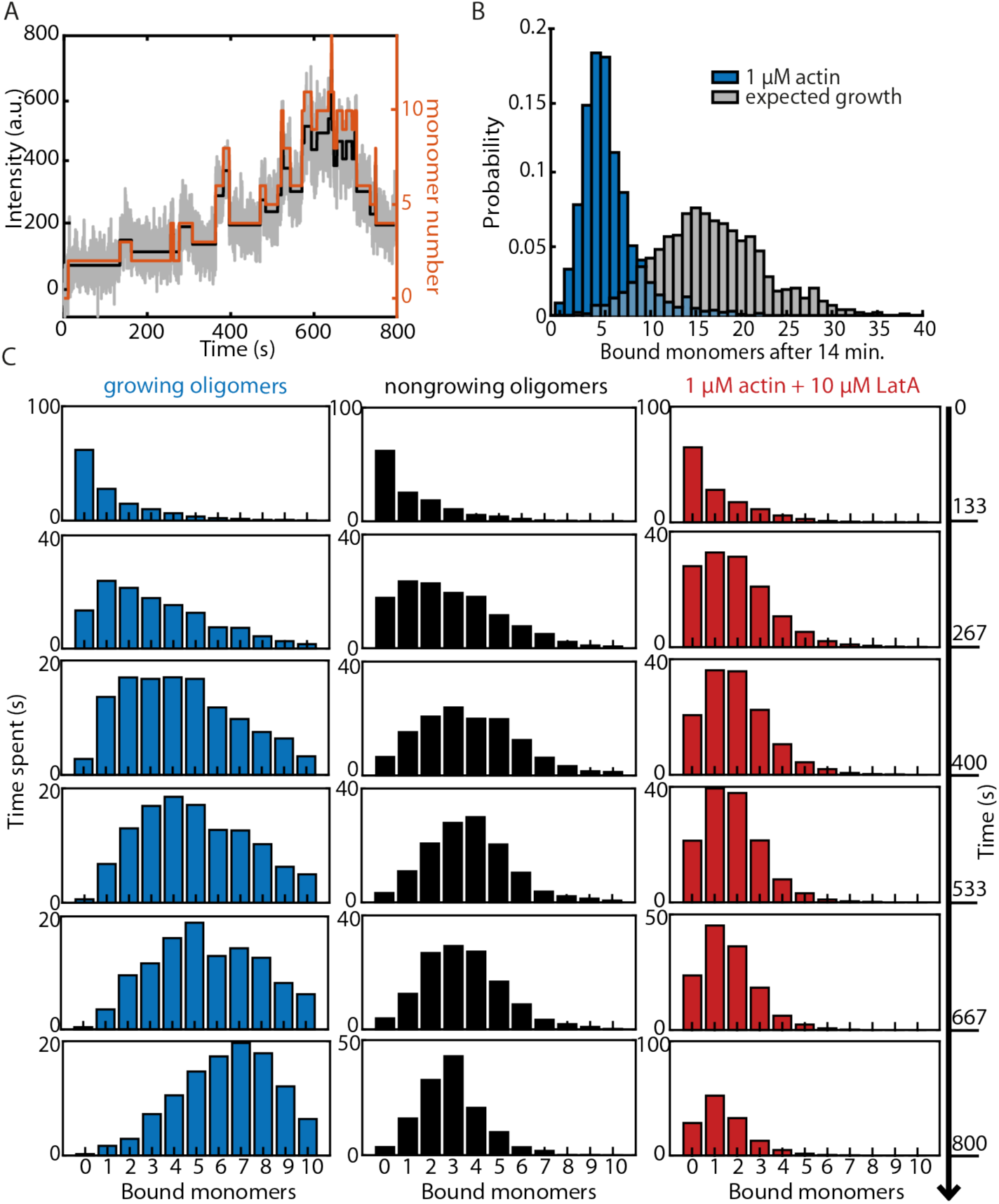
Rate extraction from intensity traces. (A) Intensity to monomer number conversion. Grey: raw time trace, black: trace as detected from step-finding algorithm, orange: conversion to monomer number. (B) Distribution of filament lengths after 14 minutes. Grey: expected behavior from a simulation using kinetic values from literature (see materials and methods for details), considering the measured bleaching rate of 0.003 s^−1^. (C) The results from a visitation analysis showing the behavior of growing and non-growing oligomers as well as the influence of LatA. The total time the oligomers spent with a certain number of oligomers was calculated for six time intervals of 133 s each.

The nucleation-elongation model of actin filament formation predicts that, upon formation of a tetramer, all formed oligomers will elongate for as long as sufficient monomers are available. In contrast to this expectation, although all individual traces reached four bound monomers at some point over the 14 min of observation, only about 10 % grew as expected (Fig. 2B). To explore in more detail the origin of growing and non-growing individual traces, we analyzed them separately, defined as traces with more than 6 or less than 4 monomers at 800 s respectively. We calculated, per 130 s time intervals, the total time a trace spent with a certain number of bound monomers. We applied the analysis to both growing and non-growing oligomers (Fig. 2C). During the first 260 s, all oligomers, i.e. filaments that will end up growing and those that will not grow, showed identical size distributions (Fig. 2C). After 260 s, the size distribution of the growing population shifted continuously over time towards larger monomer numbers (Fig. 2C), suggesting filament growth. This shift toward larger monomer numbers is independent of the N value used to define the elongating population (Fig. S11). The monomer size distribution of the non-growing oligomer population shifted its mean value to a trimer level during the first 400 s and then dwelled at that level for rest of the duration of the experiment. The observed peak in the non-growing oligomer size distribution suggests the presence of one or more rate limiting steps at the trimer or tetramer level that hinder filament formation. As a control that splitting traces into two populations does not introduce a bias in our results we repeated our analysis on simulated dynamics from a single distribution. Artificially splitting traces into two populations, when they originate from a single population, results in size histograms that are identical over time except in the last time bin used for assignment (Fig. S12). Therefore, the finding of two populations of traces represents the existence of at least two oligomer assemblies formed on the apertures.

Actin flattening, the conformational change associated with the filament structure (*3, 4*), may be involved during the initial oligomerization. To investigate the contribution of flattening during nucleation, we repeated the experiments in the presence of saturating concentrations of Latrunculin A (LatA), a macrolide that binds within the nucleotide pocket (*18*) and blocks polymerization (*19*) by preventing the inter-domain motions that allow flattening (*20*). In the presence of 10 µM LatA and 1 µM actin under polymerization conditions, traces showed on average only two bound monomers over the 800 s of experimental observation (Fig. 2C, right column). Although the maximum number of stably bound monomers was reduced in the presence of LatA, oligomers still exhibited length fluctuations (Fig. S13). Thus, LatA impairs the formation of both growing and non-growing oligomers that assemble in the absence of LatA. These results suggest that flattening of the actin molecule occurs during the dimer or trimer formation and its occurrence is necessary but does not guarantee the formation of an oligomer that can sustain growth.

Next, to gain insight into the limiting steps that forbid growth, we looked at the association and dissociation kinetics of the first six steps during oligomerization. For each trace, we recorded the time interval (‘dwell-time’) before the arrival of the next monomer (see Material and Methods for details). Dwell-time histograms were well described by single exponential functions (see Material and Methods for details) (Fig. 3 and S14). Based on this procedure, we calculated an observed association rate for each of the first six monomers for both growing and non-growing oligomers (Fig 3B). Estimated association rates revealed faster association rates for the non-growing population compared to the growing population (Fig. 3B). The difference in association rates estimated for both populations are time independent (S15), suggesting little inter-conversion between populations. The difference in association rates estimated is not a result of our sorting procedure as artificially sorting traces from the same population gives identical association rates (S16). Dissociation events cannot be accurately determined by the dwell time analysis since the loss of signal due to photo-bleaching is indistinguishable from a dissociation event. Instead, dissociation rates were estimated indirectly by comparing the experimental results with that of a Monte Carlo simulation that explicitly considers the stochastic photo-bleaching process (see Material and Methods for details). We found that only two different dissociation rates are sufficient to reproduce well our experimental observations (Fig. 3C and S17-19). Comparing the estimated dissociation rates between growing and non-growing oligomers revealed a major difference at the trimer and at the tetramer level (Fig. 3D). Trimers of growing oligomers are about a hundred times more stable than those of non-growing oligomers (Fig. 3D). At the tetramer level the scenario reverses and tetramers of non-growing oligomers are about a hundred times more stable than those of growing oligomers (Fig. 3D). A comparison of the estimated dissociation rates of the growing population and of oligomers in the presence of LatA showed the trimer level is about two orders of magnitude more stable in the growing oligomers that when flattening is blocked.

**Figure 3:**
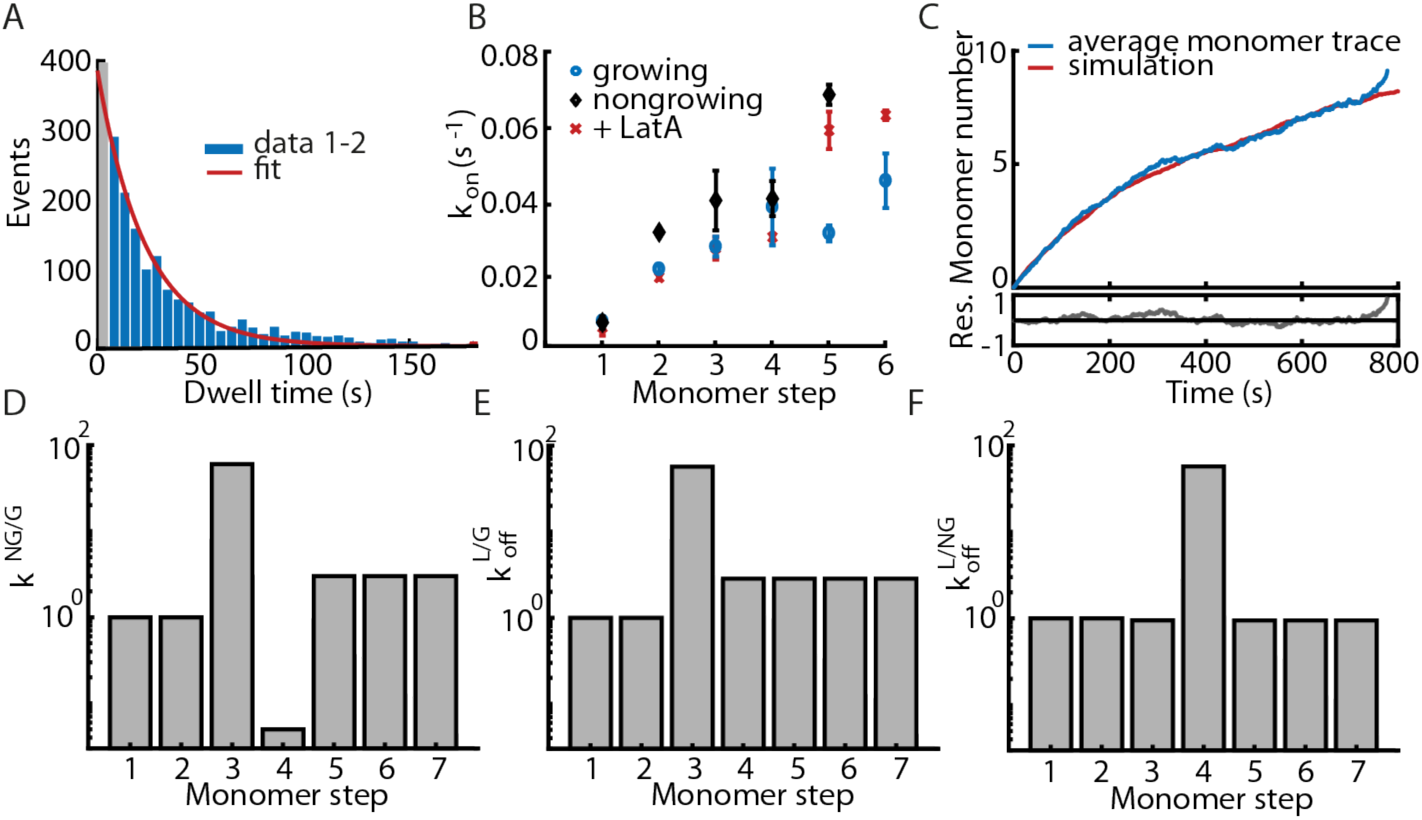
A kinetic analysis of nucleation. (A) The dwell time distribution of the second monomer-binding event. The first bin (grey) is not included in the exponential fit. (B) Association rates k_on_ for the first six binding events for 1 µM actin. Growing filaments consisted of more than six monomers (circles), nongrowing filaments of less than 4 monomers after 14 min (diamonds). (C) Mean monomer trace of the growing oligomers (blue) and simulation considering bleaching (red) for k_off_(1-3) = 0.001 s^−1^ and k_off_(>3) = 0.02 s^−1^. (D-F) Comparison of indirectly estimated off-rates for nongrowing versus growing oligomers (NG/G) (D), LatA versus nongrowing (L/NG) (E) and LatA versus growing oligomers (L/G) (F).

Furthermore, the growing oligomers showed a lower dissociation rate for all subsequent steps compared to the presence of LatA. This result suggests that flattening is required during the formation of the trimer, which stabilizes it, and it is also needed for all subsequent monomer bindings. Finally, the ratio of dissociation rates of the non-growing population and of oligomers in the presence of LatA showed that those two populations differ at the tetramer level, where the non-growing tetramers are more stable than in the presence of LatA. This result suggests that the conformational change associated with filament structure allows the formation of two different dimer arrangements.

Taking the results together, we propose that, under polymerization conditions, two dimer arrangements can be formed upon the arrival of the second monomer to a gelsolin-actin complex (Fig. 4). One arrangement, which forms faster, allows the incorporation of a third monomer although the assembled complex is unstable and growth inhibited (Fig. 4). In the other arrangement, possibly the cross-dimer (S20, see Supplemental Material for details), the addition of the third monomer is slower, the complex assembled is stable and can elongate (Fig. 4). Flattening of the actin monomer is not only required for the formation of both dimer assemblies but it is also needed for filament elongation.

**Figure 4.**
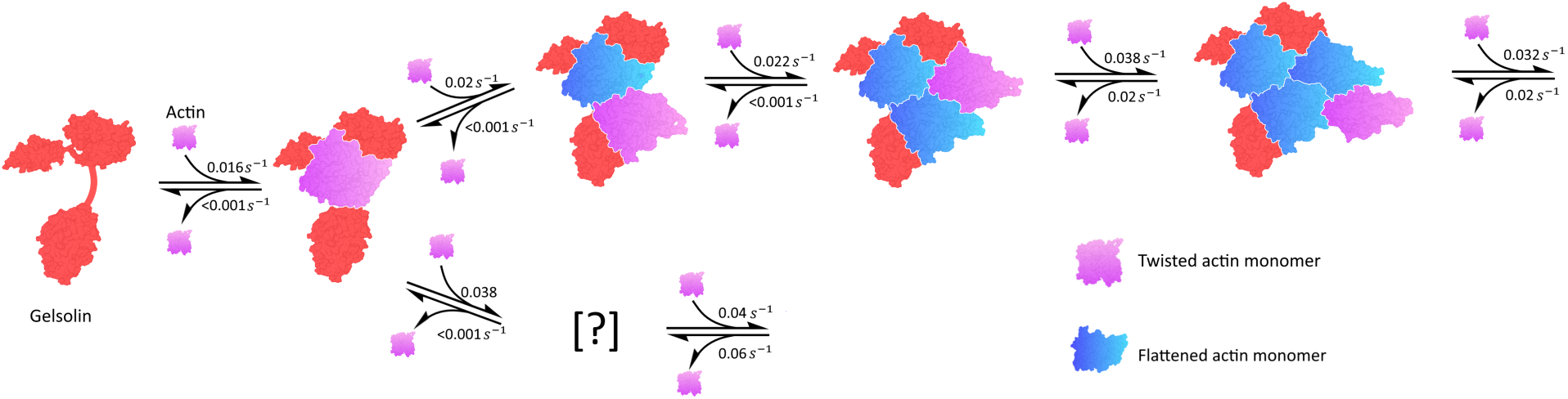
Gelsolin-mediated actin filament nucleation model. Calcium-activated Gelsolin is depicted in a ‘open’ conformation and it is colored red. Monomeric twisted actin is depicted in purple whereas flattened actin is depicted in blue. The incoming actin monomer is twisted and it flattens upon the addition of a subsequent monomer.

## DISCUSSION

Our results suggest the existence of (at least) two structurally different GA_2_ complexes. The faster association rates of this non-growing population explain how this species dominates assemblies formed. The large amount of non-growing oligomers formed during the initial steps are a possible cause of the lag phase observed in bulk experiments (*14*), since bulk assays only monitors the average polymer mass. It is possible that these non-growing oligomers represent some arrangement similar to those previously reported as AP-dimers (*21, 22*), which are known to inhibit filament formation (*21, 22*).

Our data also sheds light into the coupling between filament assembly and flattening as our data analysis suggests, *i*, that flattening occurs already during dimer formation; and *ii*, that flattening is required to stabilize the nascent filament lattice. The increase in the dissociation rate of oligomers (>4) by the presence of LatA, suggests that flattening must occur almost concomitantly with the addition of a monomer. Since flattening is associated with ATP hydrolysis (*23, 24*), this implies that mostly only the terminal monomer contains ATP (and is twisted) (Fig. 4), which agrees with previous estimates that gelsolin-capped filaments hydrolyze ATP rapidly (*25*). Moreover, the occurrence of flattening already during dimer formation may lead to ATP hydrolysis also taking place during these steps, which may be a contributing factor of the instability of dimers and trimers during filament formation of pure actin solutions.

Of the formed GA_2_ complexes that sustain elongation, we estimate that trimers are stable, with a dissociation rate upper bound at 0.001 s^−1^. The fourth and subsequent monomers dissociate with a rate of 0.02 s^−1^. Based on estimated association rates, the critical concentration of gelsolin-mediated pointed-end filament growth is about 0.5µM, in agreement with previous estimates (*16*). In contrast with the expectation from the current model of filament formation that predicts a dissociation rate of the third monomer to be two to three orders of magnitude larger than that of subsequent additions, we estimate that trimers that sustain elongation are stable. Therefore, a competing pathway (Fig. 4) and not thermodynamically unstable oligomers determine gelsolin-actin filament formation. The extent of biasing dimer arrangement towards those that can sustain elongation may be the mechanistic difference between strong and weak actin filament nucleators. The use of ZMW to directly observe the first steps during monomer assemble will allow answering those questions not only for actin but also for any other filament system.

## Material and Methods

### Proteins and DNA

Atto488-labeled actin and biotinylated gelsolin were purchased from Hypermol (Bielefeld, Germany). We have shown that fluorescently labeled actin is fully active as assayed by bulk techniques, TIRF and single molecule methods (*26*). The capping behavior of biotinylated gelsolin was not impaired as compared to unlabeled gelsolin (Fig. S2). Labeled DNA double strands were obtained from IBA (Göttingen, Germany). DNA molecules were labeled with 4, 6, or 8 atto488 dyes and attached to a biotin-linker.

### Experiments

Zero Mode Waveguides (ZMWs, Pacific Biosciences, Menlo Park, Canada) were incubated with 0.15 mg/ml Streptavidin and 5 mg/ml BSA in PBS for 5 min. After washing with PBS, 200 nM freshly diluted biotinylated gelsolin was incubated for 5 min, followed by 5 min incubation with 1 mg/ml BSA and 5 mg/ml biotinylated BSA. Actin-atto488 in G-buffer was incubated in magnesium exchange buffer for 5 minutes on ice (50 µM MgCl_2_, 0.2 mM EGTA). Latrunculin A (LatA) at 10x the actin concentration was added after 4 minutes and incubated for 1 minute. Actin polymerization was induced by adding 1/10 volume of a 10x concentrated polymerization buffer (50mM KCl, 2 mM MgCl_2_, 0.2 mM CaCl_2_, 0.2 mM ATP, pH 7.0). The final buffer contained a PCA/PCD oxygen scavenging system with 250 nM protocatechuate dioxygenase ‘PCD’, 2.5 mM 3,4-dihydroxybenzoic acid ‘PCA’ and 1 mM Trolox (*27*). After starting polymerization, 50 µl of the reaction mixture were added onto the waveguides and data acquisition was started immediately. Experiments were carried out a home-built wide field microscope system using a 60x 1.45 NA oil immersion objective (Plan Apo TIRF 60x, Nikon). The laser power of a 488 nm laser (Cobolt MLD, Cobolt AB) was set to 5 mW before entering the objective via a dichroic mirror (zt405/488/561/640rpc, AHF Analysentechnik). After passing through an emission filter (535/50 BrightLineHC, AHF Analysentechnik), the signal was recorded by an EMCCD camera (Andor iXon, Andor Technology). The intensity as a function of time per aperture was extracted using custom written software in MATLAB (The MathWorks). To measure the bleaching rate, 1 µl of Phalloidin was added to the waveguides after 1.5 h to stabilize the formed filaments and prevent dissociation.

DNA measurements were carried out using 100 nM DNA, labeled with 4, 6 or 8 atto488 dyes in PBS. DNA was added to the waveguides after streptavidin incubation as described above. After incubation for 30 min and washing with PBS, acquisition was started.

Fluorescently labeled actin was also assayed in TIRF for functionality and to control that biotinylation of gelsolin did not affect its behavior. Actin filaments were imaged in TIRF as reported previously (*28*). To measure the gelsolin capping behavior, before mixing with polymerization buffer, 5 nM gelsolin or biotinylated gelsolin was added and incubated for 1 min.

### Data analysis

#### The step-finding algorithm

To estimate the position in time and the change in intensity associated with monomer addition or dissociation, we used a recently developed algorithm (*17*). The algorithm builds a cost function with the proposed steps and the noise present. To prevent overfitting the algorithm uses a penalty factor for the introduction of new steps (*17*). We compared the results obtained by our method with those estimated by combining a L1 regularization method (*29*) with a model-independent step finding algorithm (*30*), as has been used previously for optical tweezers data (*31*). We tested the step-finding algorithms on simulated data with a signal-to-noise ration (SNR) of 1 (Fig. S3). The number of steps in a trace and the step size distribution reflected the input parameters. To test the analysis on experimental data, we used DNA double strands labeled with 4, 6 or 8 attto488-dyes. The number of down-steps obtained from bleaching experiments reflected the number of dyes on the double-stranded DNA (Fig. S4). The Salapaka algorithm (*17*) performed faster and had a smaller error in detection of the number of steps (Fig. S3).

#### Conversion of intensity traces to oligomer length

After the position and size of each step-wise transition was assigned, we converted the observed intensity levels into a monomer number. The conversion process followed the following steps: we first estimated the mean step size per experimental condition by fitting the step size distribution with a gamma function. Then, using the estimated mean-step size, we binned the intensity levels and assigned all events within that bin to the same monomer number. We also observed intensity transitions that were significantly higher than twice the mean step size. These transitions may represent sequential binding reactions that occur too fast to be detected individually. To correct for those, we split them into 2 or 3 individual steps. The dwell time of the newly introduced steps is assigned to be equal to the frame rate of 200 ms. Therefore, it will appear in the first bin of the dwell time distribution, which is excluded in the single-exponential fit (Fig. S12). With the double-step correction, we obtain reproducible step-size distributions for all measurements (Fig. S5).

#### Association Rates

Association rates were determined from a single-exponential fit of the dwell time distributions. The dwell times are defined as the total time spent on monomer number *n* until the next monomer *n*+1 arrives. Only events before the first down step were considered, since down events cannot be distinguished between dissociation (that changes the monomer number) and photo-bleaching (that does not change the monomer number).

To estimate the total time an oligomer spends at a particular number of monomers, we divided the 800 s traces into 6 time intervals and the time spent in each monomer level was calculated.

#### Simulation of nucleation

Simulations of a stepping process with 7 different on- and off-rates per step (6 individual monomer transitions plus elongation) were performed with 1000 cycles per concentration. The resulting traces were overlaid with Gaussian noise with a SNR of 1. Simulated traces were analyzed in the same manner as experimentally obtained traces.

#### Dissociation Rates

We used a model with 7 different on- and off-rates per step (6 individual monomer transitions plus elongation) and compared the average trace of 2000 simulated runs with the average of the experimental traces (Fig. S15-S17). We calculated the χ^2^ between the simulated average trace and the experimentally obtained. We sampled each individual off rate. While doing this we realized that only two different rates were needed to describe the experimental average trace.

#### Previously reported values

We used the following reference values from the literature: Binding of the first actin monomer to gelsolin is 0.025 µM^−1^∙s^−1^ at pH 7.0 (*32*) (0.015µM^−1^∙s^−1^ (4)). Binding of second actin monomer to gelsolin-actin complex is ~40 times faster than binding of the first actin monomer (0.8 µM^−1^∙s^−1^) (3). Estimated pointed-end *k*_on_ from Gelsolin-(actin)_2_ is 0.12 µM^−1^∙s^−1^ (*33*). Dissociation of the second actin monomer (i.e. Gelsolin-(actin)_2_ to Gelsolin-(actin)_1_) is 0.02 s^−1^ (*34*). Gelsolin-mediated pointed-end *k*_on_ 0.5 µM^−1^∙s^−1^, *k*_off_ 0.32s^−1^ (*35*).

Pointed end *k*_on_ is 1.3µM^−1^∙s^−1^, *k*_off_ is 0.8 s^−1^ (*16*).

### Structural model

A structural arrangement was generated out using Chimera (*36*). Since there is no structure of full-length gelsolin with actin we used available structures of gelsolin domains G1-G3 (1RGI (*37*)) and G4-G6 (1H1V (*38*)) with actin and Ca^+2^. We then aligned the actin of the G1-G3 gelsolin-actin complex to the terminal actin monomer within the filament structure (*39*) (monomer 1 in Fig. S18). The actin of the G4-G6 gelsolin-actin complex was then aligned to the monomer on the opposite strand within the filament (monomer 2 in Fig. S18). This arrangement corresponds to a cross-dimer configuration and has been proposed before (*40*). The calculated the distance between the C-terminal amino acid of the G1-G3 chain and the N-terminal amino acid of the G4-G6 fragment is about 6 nm, which is too long to be traversed by the missing linker of about 40 amino acids. Since no other arrangement generated shorter distances (data not shown), this suggests that further conformational changes take place during gelsolin-actin complex formation. The structure of the full-length gelsolin with the actin filament end remains to be addressed.

## Acknowledgements

This work was supported by a grant from the Deutsche Forschungsgemeinschaft through the SPP 1464 (to AHC and DCL), the Excellent Clusters Nanosystems Initative Munich (NIM) and Center for Integrated Protein Science Munich (CIPSM), and the Ludwig-Maximilians-Universität München (via the LMUInnovativ BioImaging Network (BIN) and the Center for NanoScience (CeNS)). Work in the Crevenna laboratory was financially supported by: Project LISBOA-01-0145-FEDER-007660 (Microbiologia Molecular, Estrutural e Celular) funded by FEDER funds through COMPETE2020 - Programa Operacional Competitividade e Internacionalização (POCI) and by national funds through FCT - Fundação para a Ciência e a Tecnologia.

---

**Figure S1:**
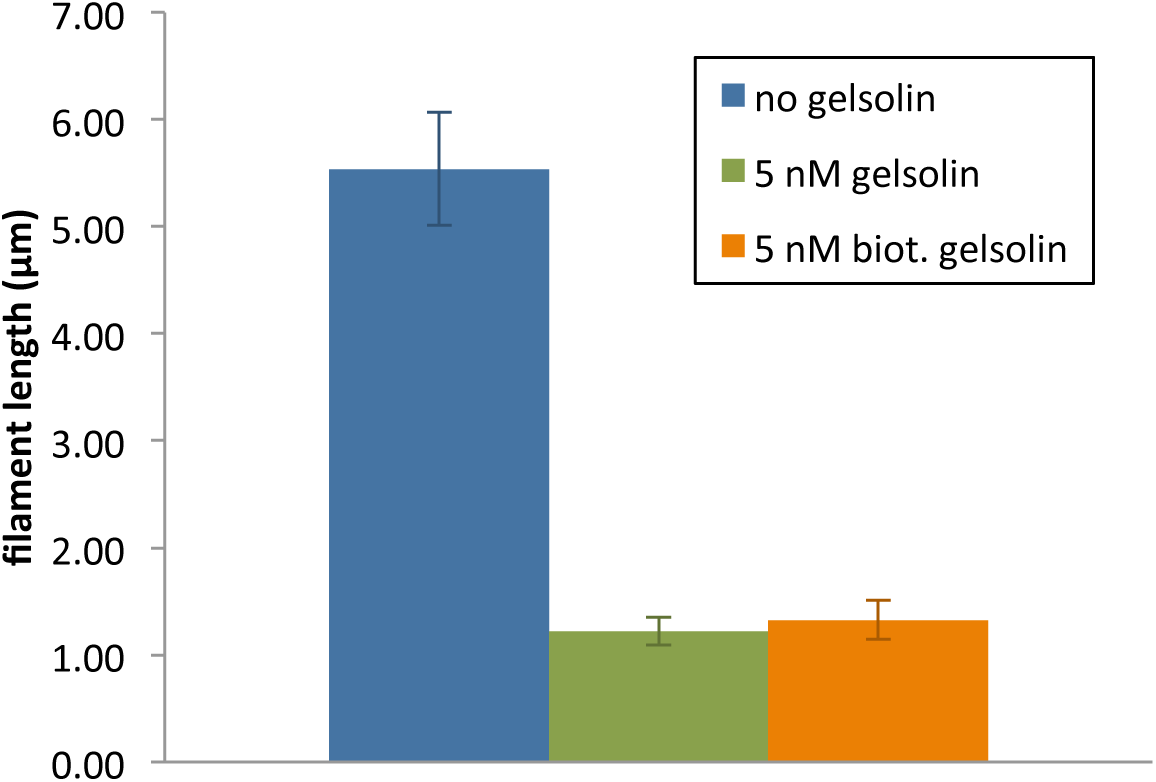
Biotinylation of gelsolin does not impair capping. Mean filament length after 5 minutes growth of 1 µM actin and 5 nM gelsolin or biotinylated gelsolin with a degree of labeling of 0.8.

**Figure S2:**
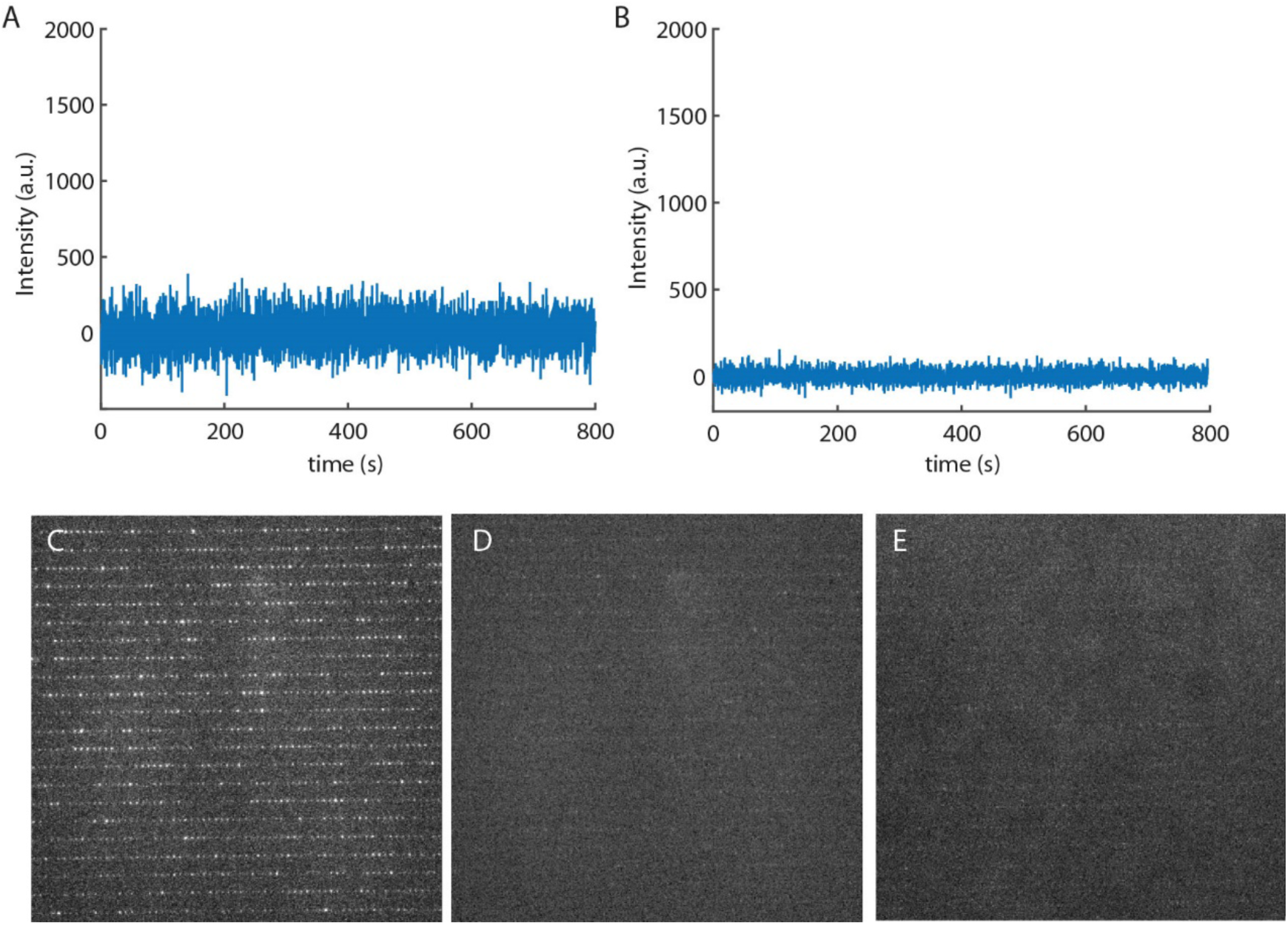
Intensity changes signifying nucleation were not observed in G-Buffer or without gelsolin. (A) An example trace from a ZMW for a measurement in G-Buffer. (B) An example trace from a ZMW for a measurement without gelsolin as a tethering protein. (C) An image of a waveguide array with 200 nM gelsolin and 1 µM actin in polymerizing conditions after 14 minutes. (D) An image of a waveguide array with 200 nM gelsolin and 1 µM actin in G-Buffer after 14 minutes. (E) An image of a waveguide array with 1 µM actin without gelsolin in polymerizing conditions after 14 minutes.

**Figure S3:**
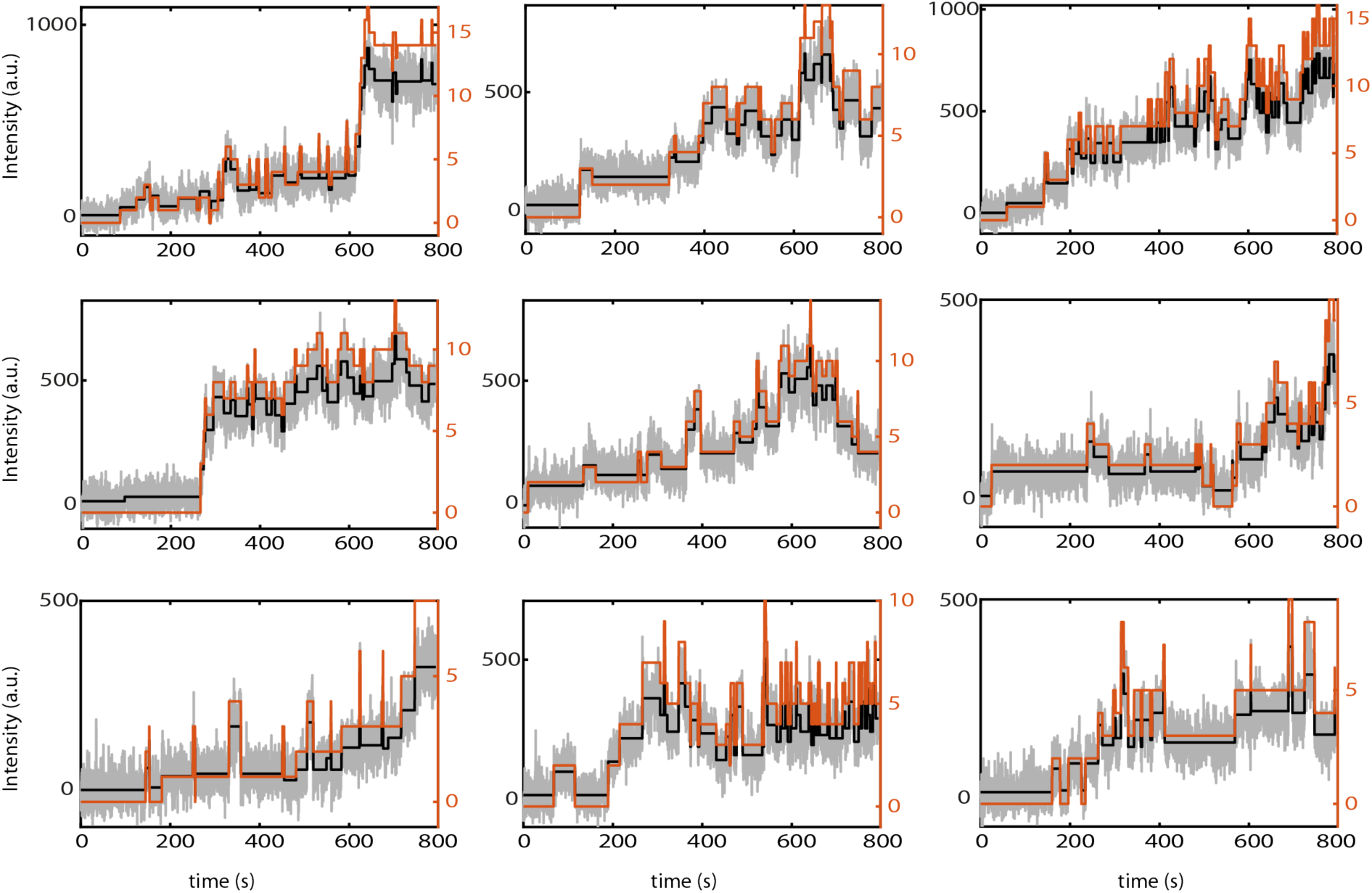
Intensity to monomer conversion. A few random traces are shown for 1 µM actin on gelsolin. Grey: fluorescence intensity, black: result of the step-finding algorithm, orange: conversion to monomer number, using the step-size information and double-step correction.

**Figure S4:**
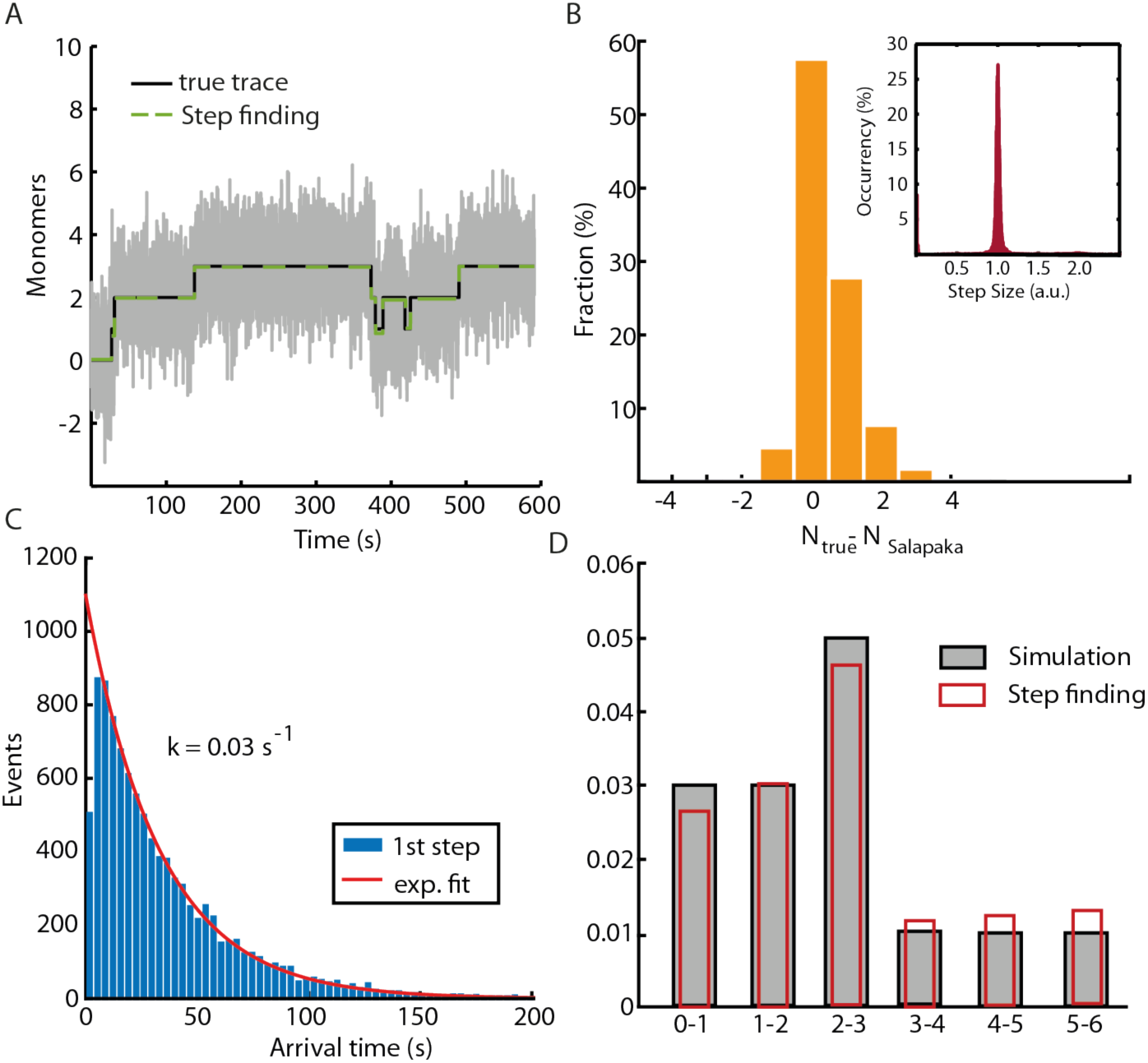
Testing of the step-finding algorithm. (A) A simulated trace (black) with overlaid noise (SNR = 1) (grey) and the result of the Salapaka step-finding algorithm (dashed green line). (B) Discrepancy between the true simulated number of steps in a trace and the number of steps as found by the step-finding algorithm. Inlets: Step size distribution. (C) Arrival time distribution for the first up-step with single-exponential fit (red line). (D) On-rates obtained from the exponential fits of the arrival times for the different steps. Grey: simulated input rates, red: recovered rates from the noise-overlaid simulations.

**Figure S5:**
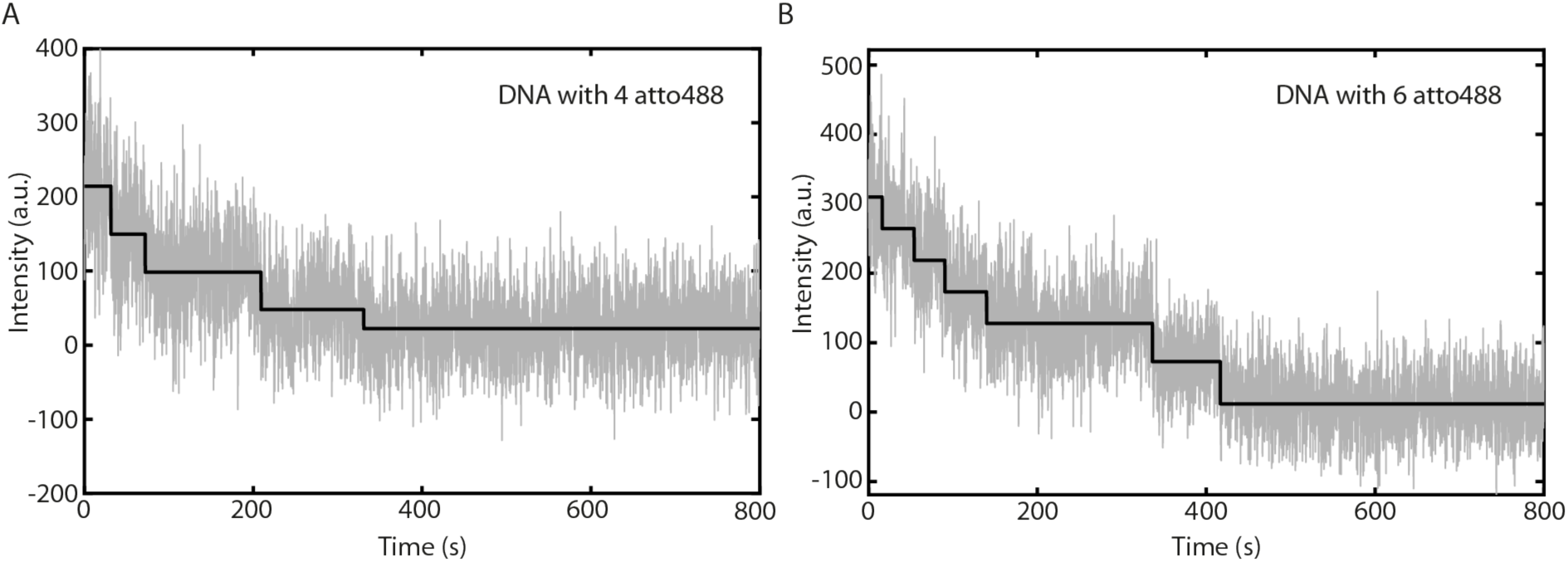
Example of a bleaching trace of a DNA double strand labeled with 4 atto488-dyes. Grey: measured fluorescence signal, black: result of the step-finding algorithm.

**Tab. 1:** Mean number of extracted down steps for DNA strands with 4, 6, and 8 atto488-dyes, respectively. For step-finding, the Salapaka algorithm was used with a penalty factor of 50.

| 4 dyes | 6 dyes | 8 dyes |
| --- | --- | --- |
| $5.5 \pm 1.3$ | $6.7 \pm 1.3$ | $7.3 \pm 1.4$ |

**Figure S6:**
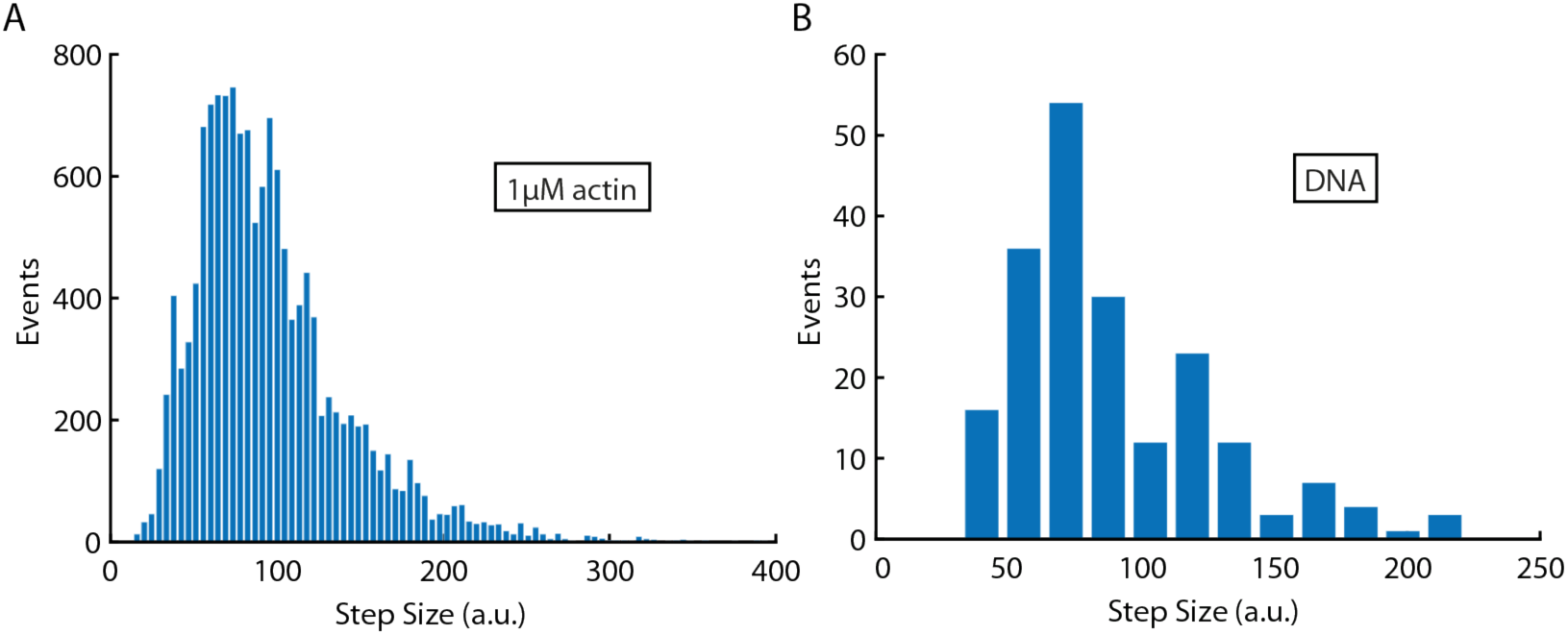
Step sizes are reproducible among measurements. The step size distribution determined for 1 µM actin (A) and DNA-atto488 (B), respectively, are shown.

**Figure S7:**
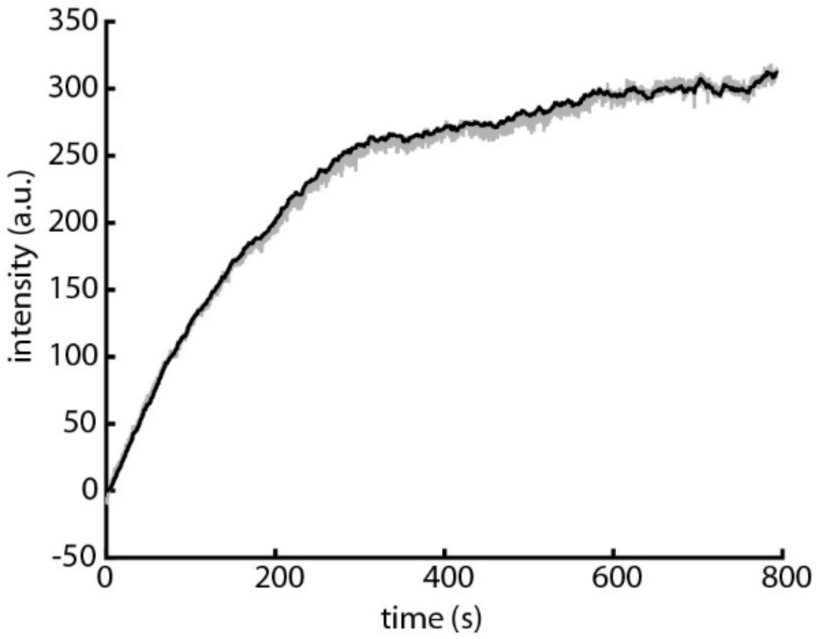
Intensity to monomer conversion. Mean intensity trace for 1 µM actin (grey, mean of about 600 traces) and extracted monomer trace, multiplied by the mean step size (black).

**Figure S8:**
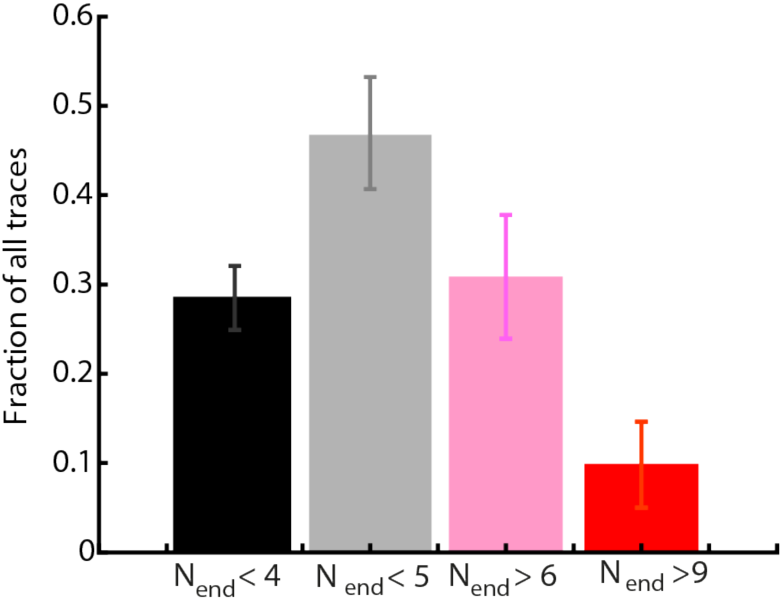
Fraction of oligomers that show less than 4 or 5, or more than 6 or 9 monomers after 800 s.

**Figure S9:**
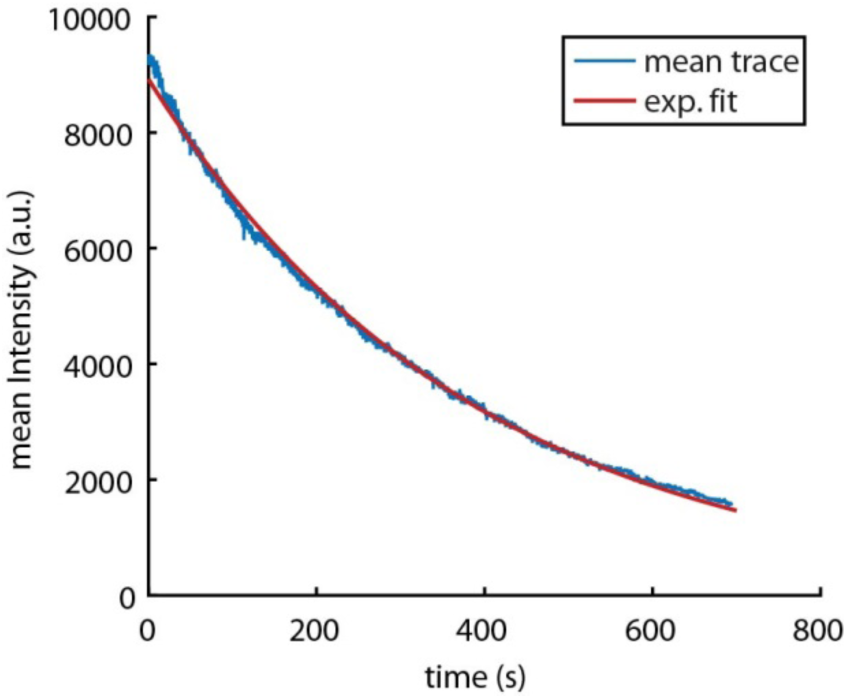
Bleaching rate of stabilized filaments. 1 µM actin was incubated for 1.5 h and stabilized with phalloidin to prevent dissociation. Blue: mean fluorescence intensity decay from about 600 traces, red: exponential fit with decay constant k = 0.003 s^−1^.

**Figure S10:**
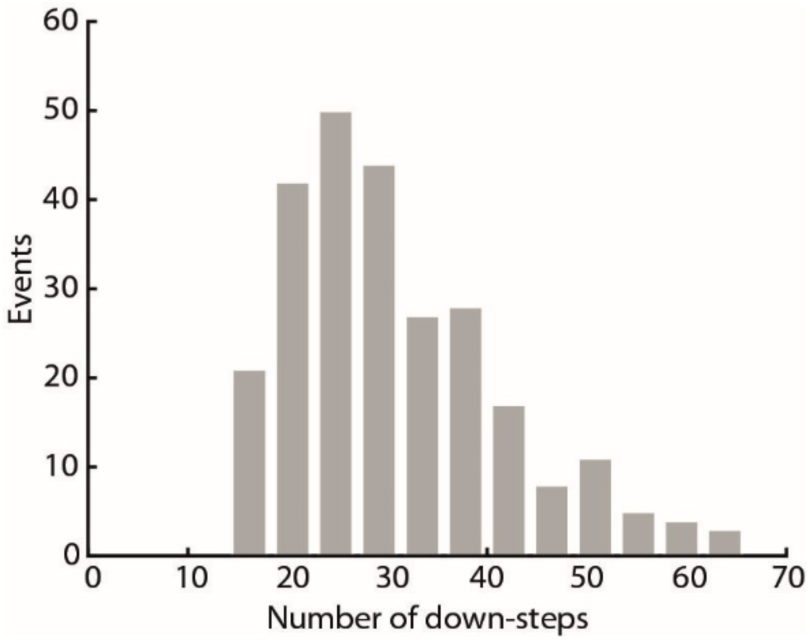
Long filaments can be observed in the waveguides. Histogram of the number of steps during bleaching of 1 µM actin polymerized on surface-immobilized biotinylated actin after 1 hour incubation in the ZMWs.

**Figure S11:**
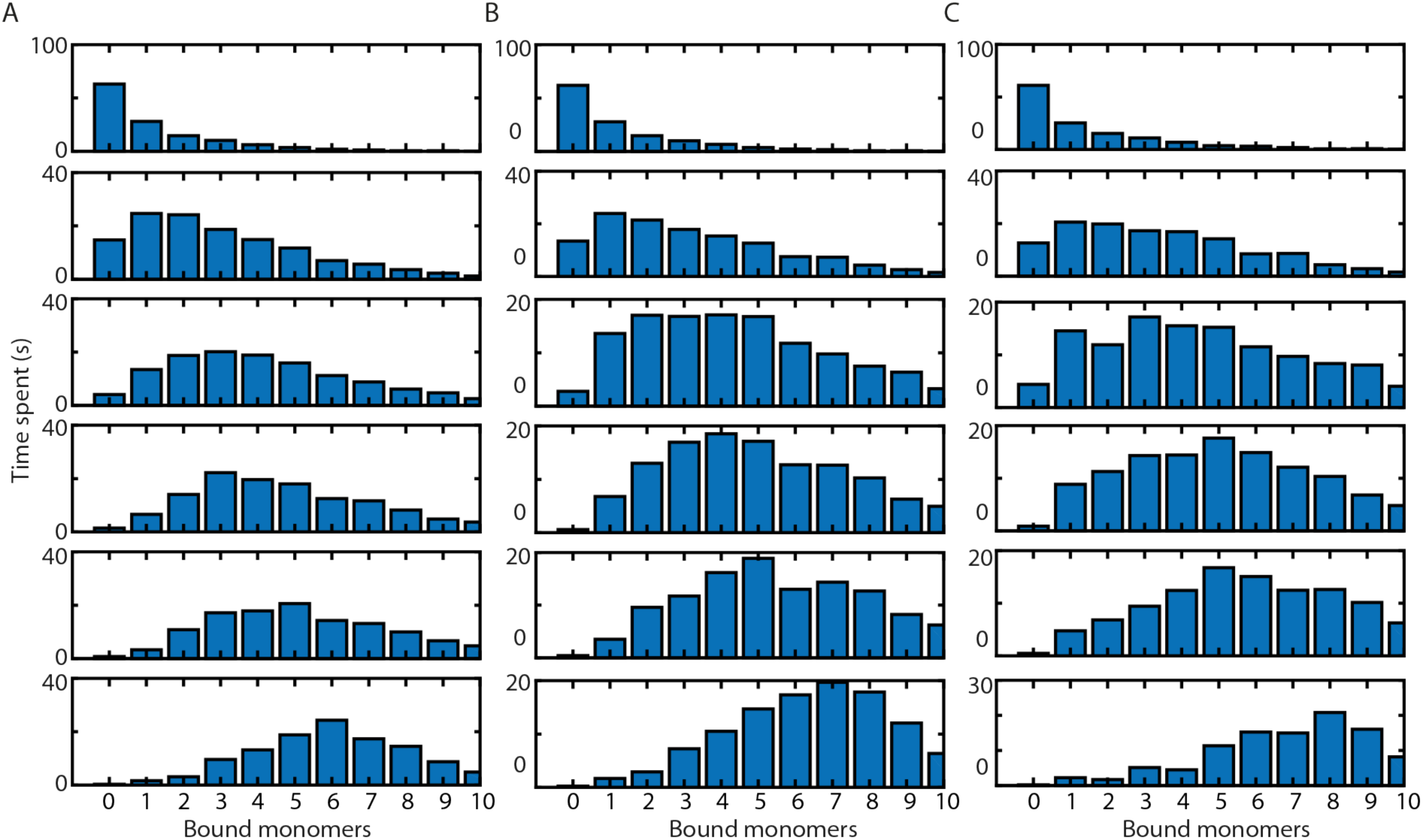
Visitation analysis on growing oligomers that contain of five monomers (A), six monomers (B) or seven monomers (D).

**Figure S12:**
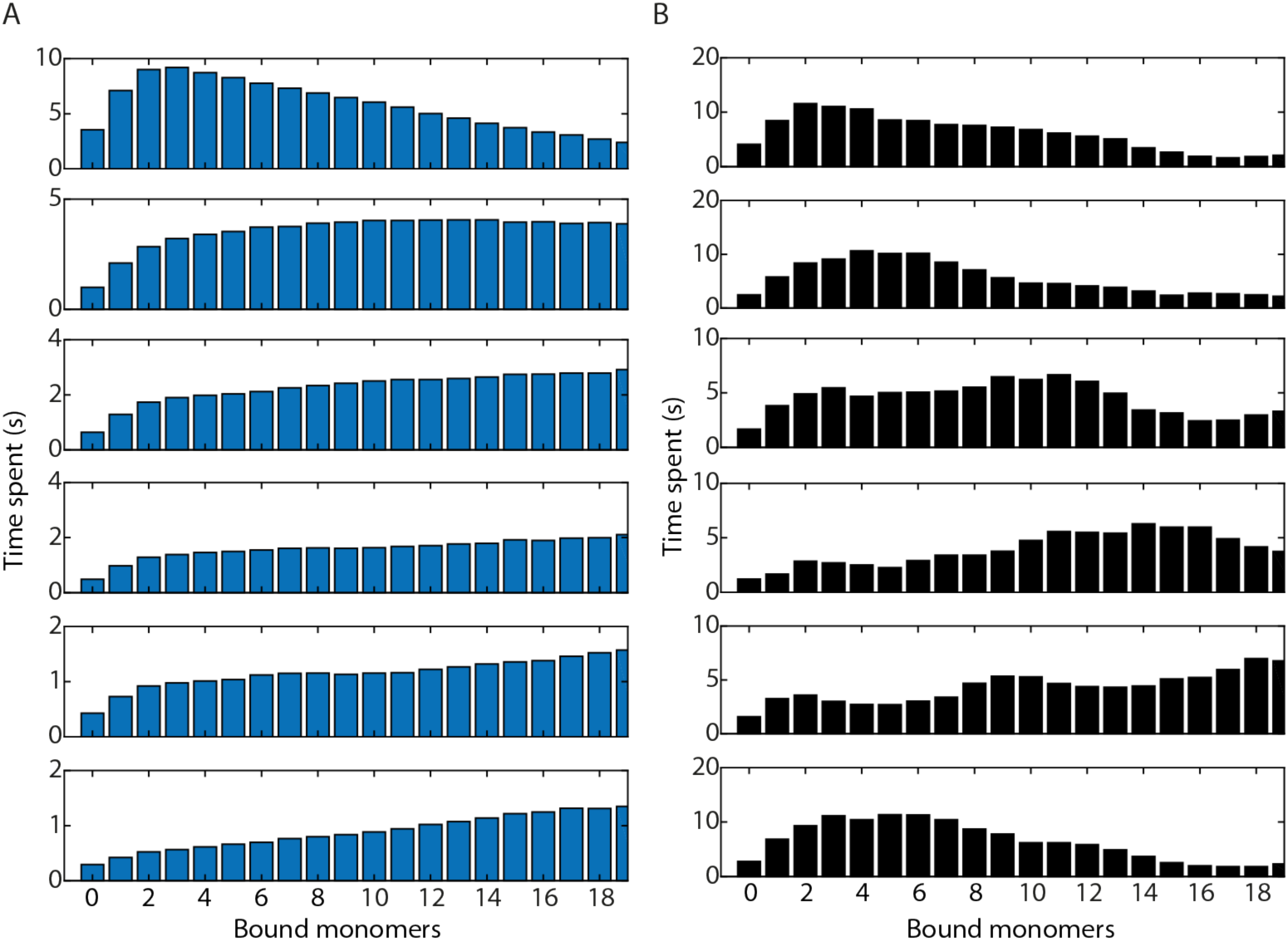
Visitation analysis on a simulation of a random walk with an association rate of 1 s^−1^ and a dissociation rate of 0.95s^−1^ (see materials and methods for details). The simulated traces were divided into a “growing” (A) and a “non-growing” (B) population based on the monomer number after 800 s. “Non-growing” show below 4 monomers after 800 s, “growing” above 6 monomers.

**Figure S13:**
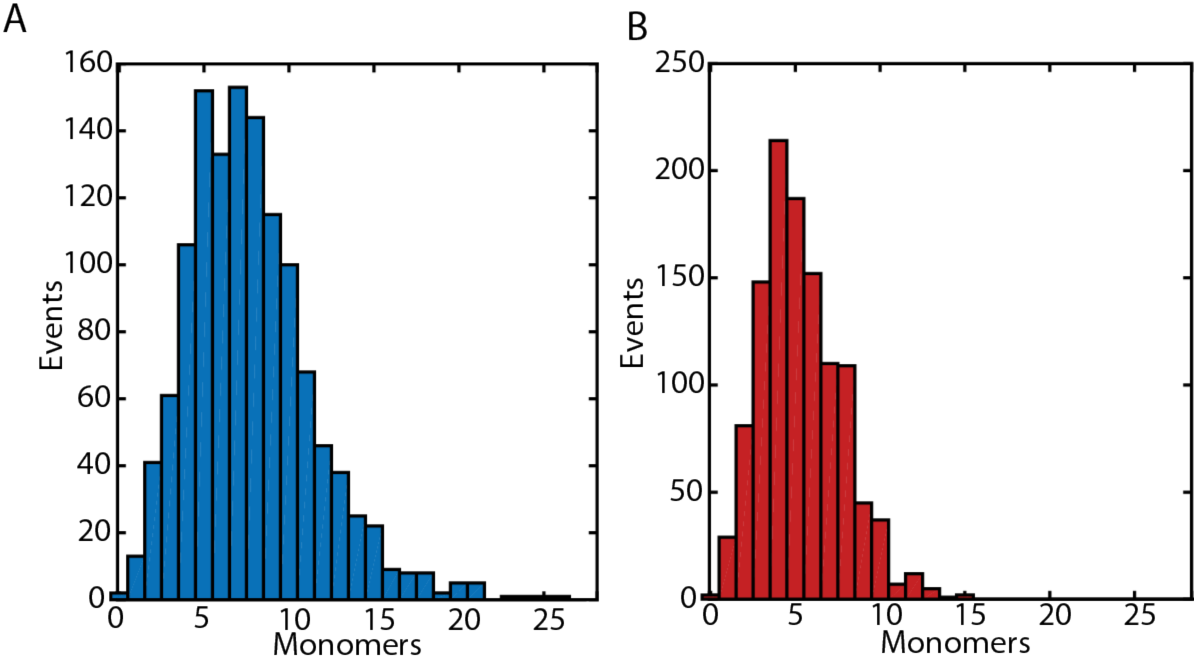
The maximum number of bound monomers during an individual trace. (A) 1 µM actin, (B) 1 µM actin with 10 µM LatA.

**Figure S14:**
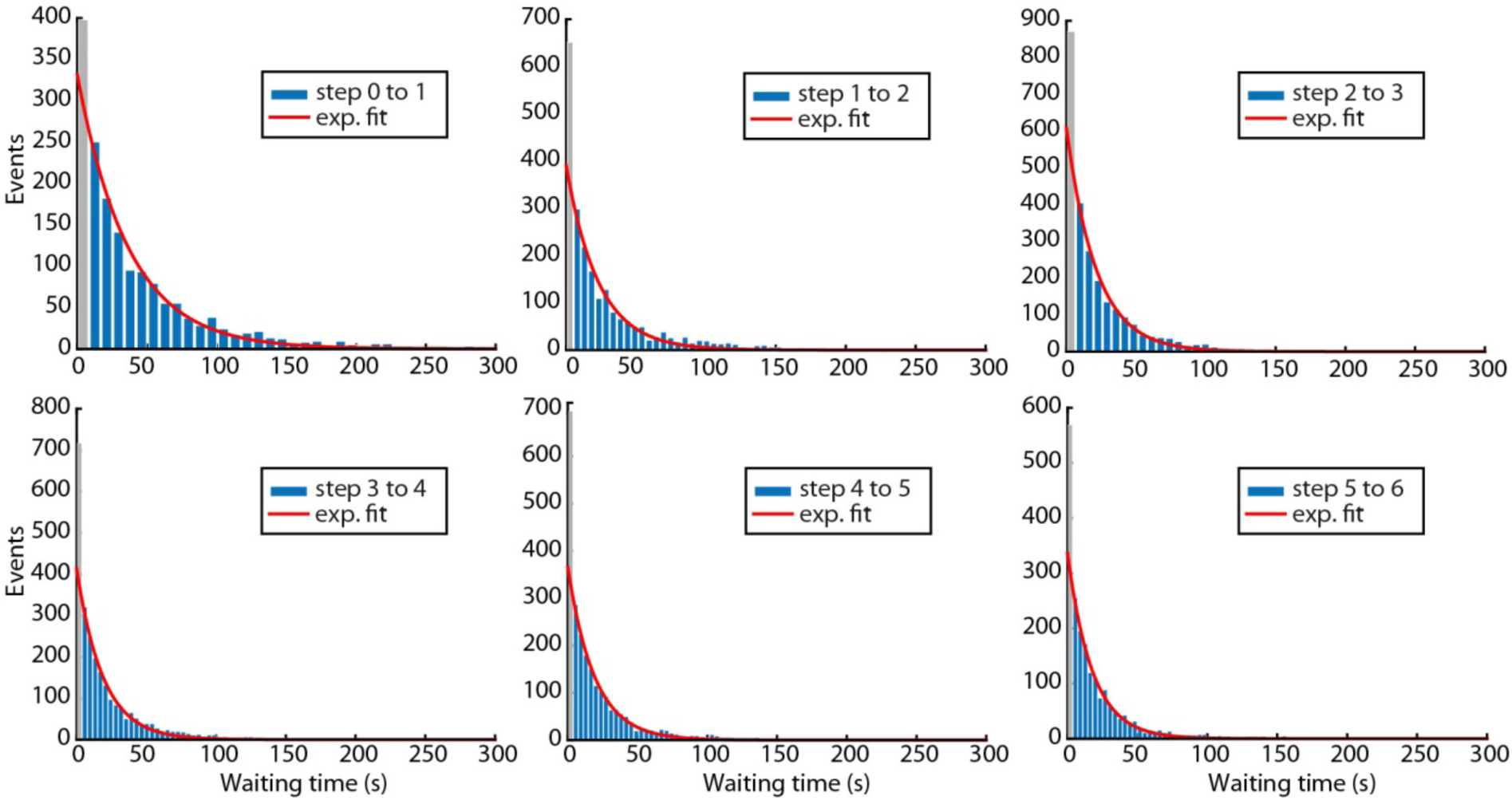
Association rates are single-exponentially distributed. Waiting times (blue) and exponential fits (red) of the first to the sixth binding events. The first bins (grey) were not included in the fits.

**Figure S15:**
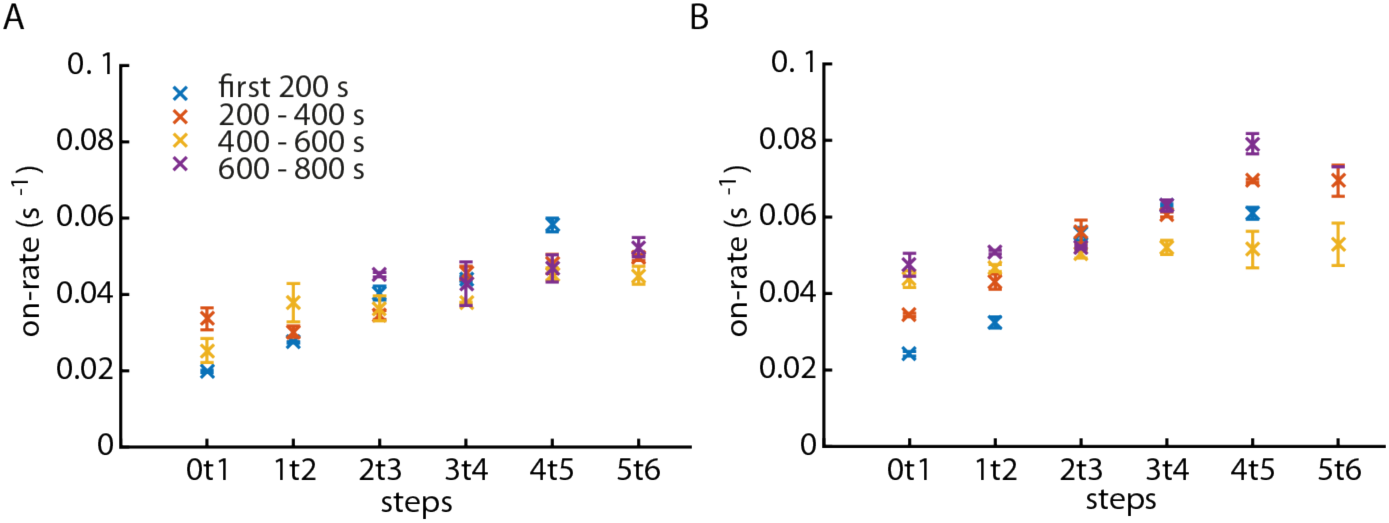
Association rates do not change over time. The time course of the measurement was divided into four time intervals of 200 s each. For each time interval, the association rates of single steps were determined independently for (A) growing and (B) non-growing oligomers. Data points with insufficient statistics have been omitted.

**Figure S16:**
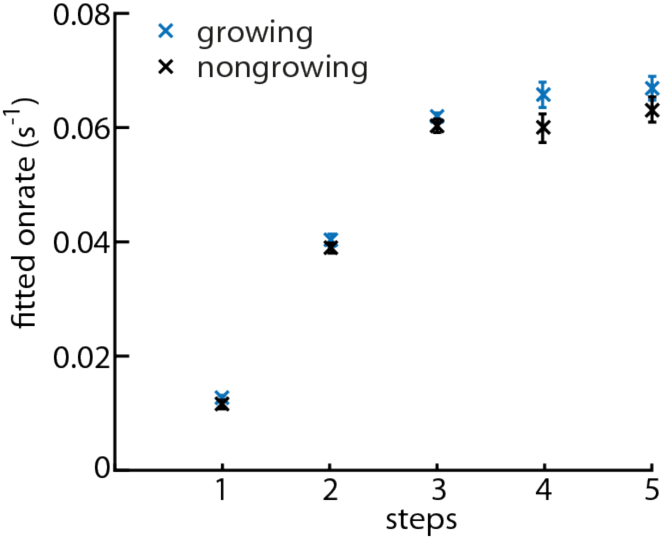
Association rates determined from a simulation of literature values (see materials and methods for details). The simulated traces were divided into a “growing” (blue) and a “nongrowing” (black) species based on the monomer number after 800 s. The “nongrowing” species showed less than 4 monomers after 800 s, the “growing” species contained above 6 monomers.

**Figure S17:**
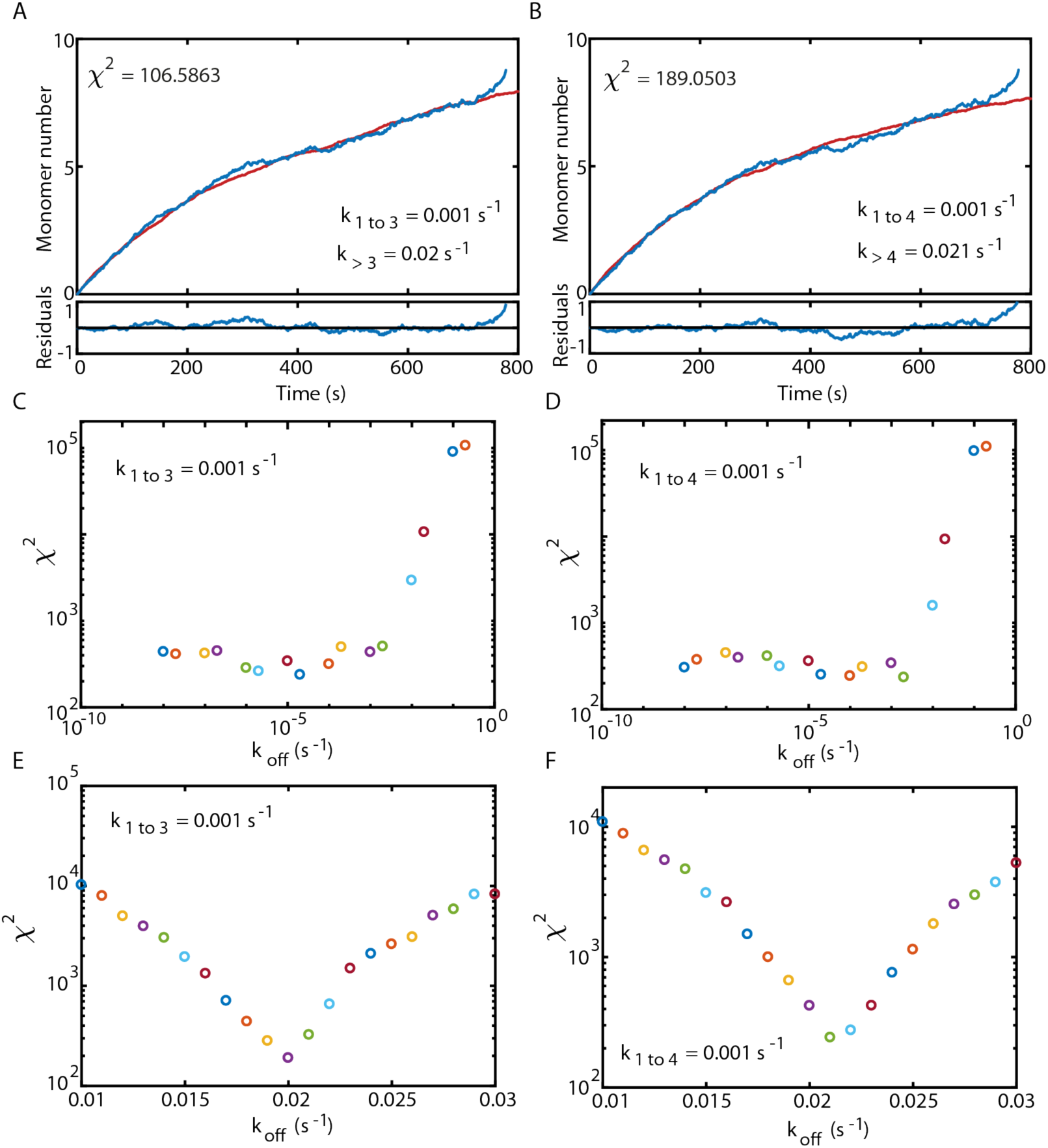
Offrate estimation for growing oligomers. (A, B) Comparison between the average of all growing oligomer traces (blue) and a simulation using the measured association rates and two different dissociation rates (red). The dissocation rate for the first three (A) or four (B) monomers was 0.001 s^−1^, for each subsequent step 0.02 s^−1^ or 0.021 s^−1^. (C-F) Chi^2^ error for the comparison between experiment and simulation for a dissociation rate of 0.001 s^−1^ for the first three (C) or four (D) steps for different dissociation rates for the subsequent steps. (E,F) For both models, a minimum in the Chi^2^ error can be reached.

**Figure S18:**
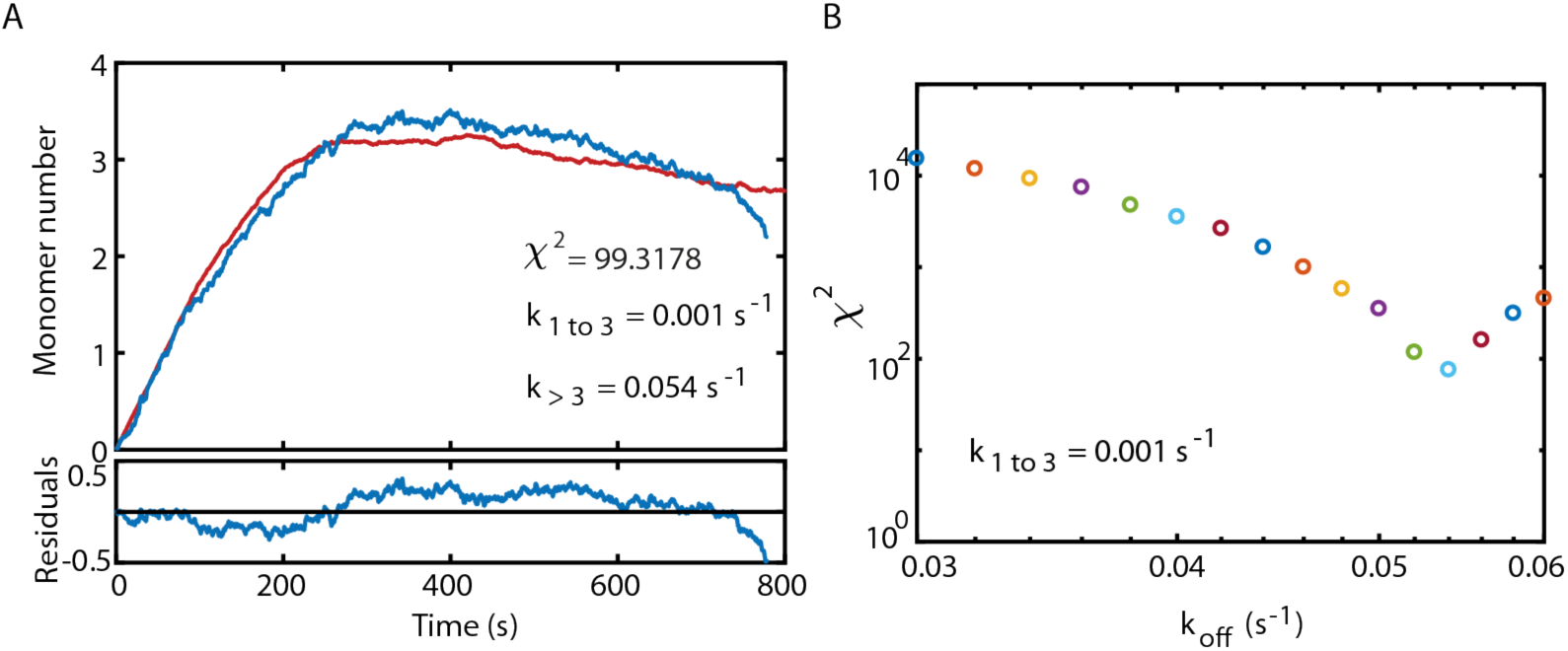
Offrate estimation for non-growing oligomers. (A) Comparison between the average of all growing oligomer traces (blue) and a simulation using the measured association rates and two different dissociation rates (red). The dissocation rate for the first three monomers was 0.001 s^−1^, for each subsequent step 0.054 s^−1^. (B) Chi^2^ error for the comparison between experiment and simulation for a dissociation rate of 0.001 s^−1^ for the first three steps for different dissociation rates for the subsequent steps.

**Figure S19:**
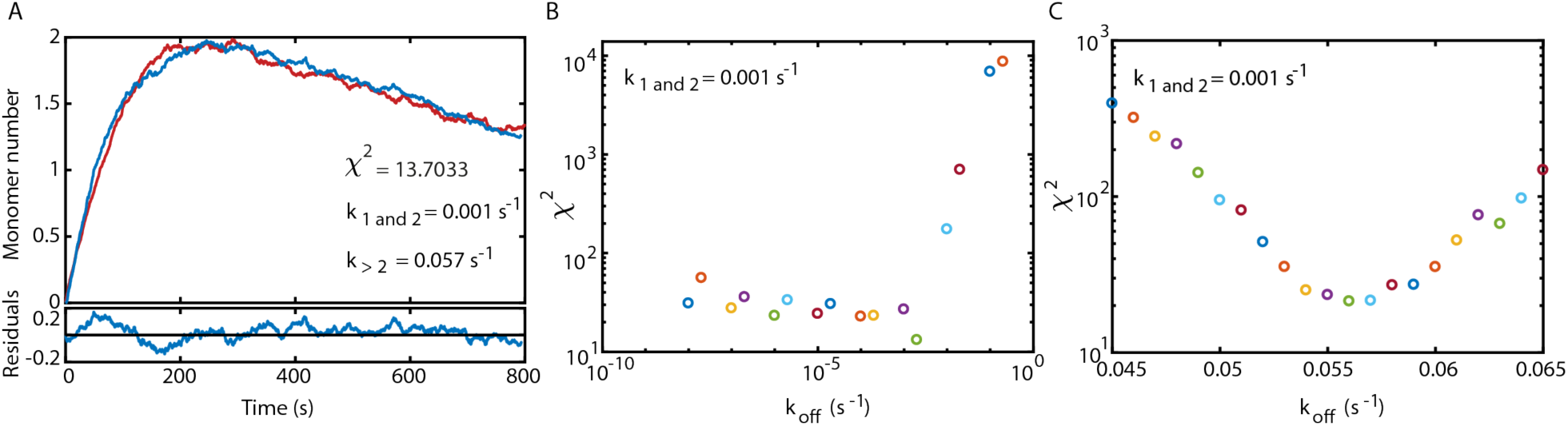
Offrate estimation for oligomers with LatA. (A) Comparison between the average of all growing oligomer traces (blue) and a simulation using the measured association rates and two different dissociation rates (red). The dissocation rate for the first three monomers was 0.001 s^−1^, for each subsequent step 0.057 s^−1^. (B, C) Chi^2^ error for the comparison between experiment and simulation for a dissociation rate of 0.001 s^−1^ for the first three steps for different dissociation rates for the subsequent steps.

**Figure S20:**
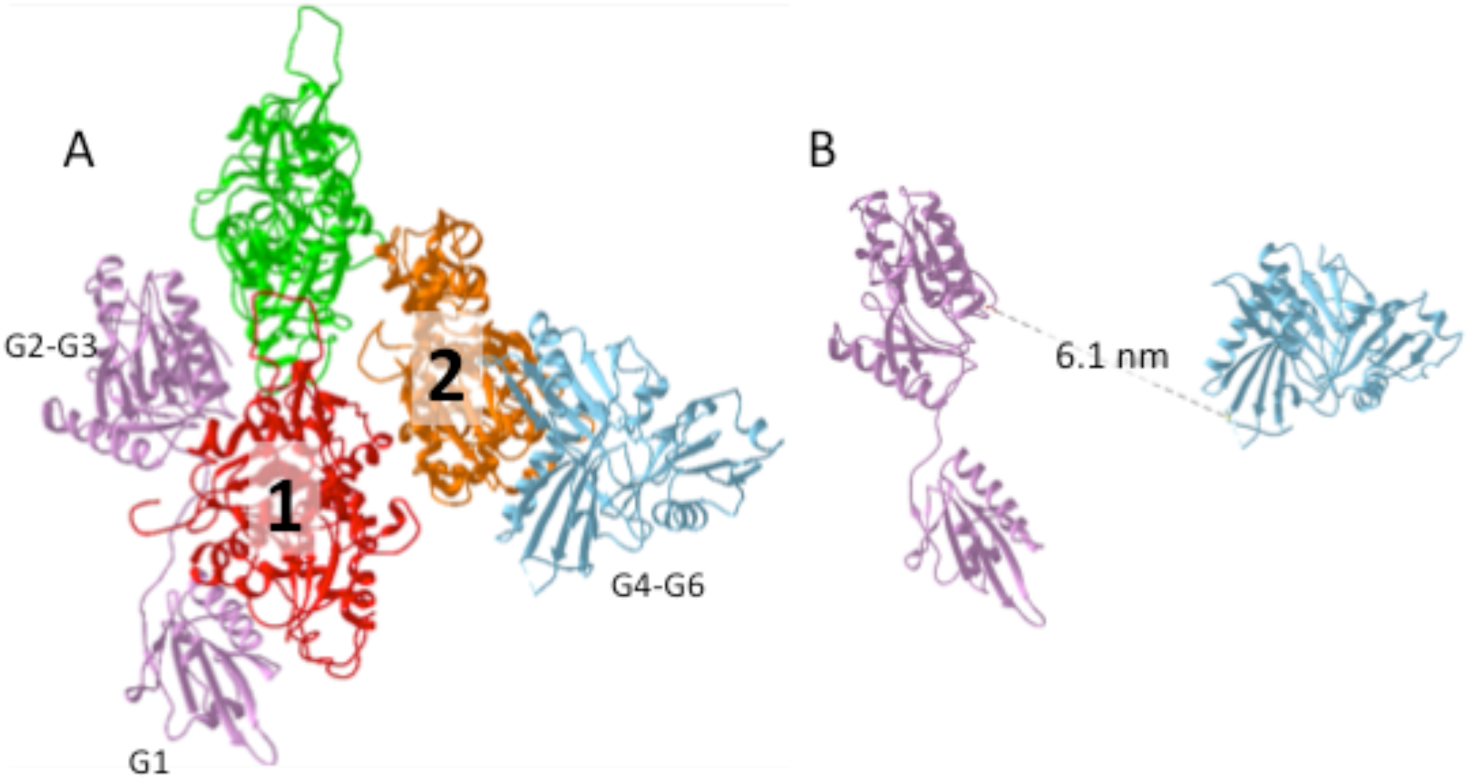
Structural model of full-length gelsolin bound to the actin filament barbed-end. (A) Alignment of G1-G3 (1RGI) and G4-G6 (1H1V) gelsolin fragments complexed with actin to the filament barbed-end. (B) measured distance between the C-terminal amino acid of G3 to the N-terminal amino acid of G4.

